# Establishment of locally adapted mutations under divergent selection

**DOI:** 10.1101/248013

**Authors:** Matteo Tomasini, Stephan Peischl

## Abstract

We study the establishment probabilities of locally adapted mutations using a multitype branching process framework. We find a surprisingly simple and intuitive analytical approximation for the establishment probabilities in a symmetric two-deme model under the assumption of weak (positive) selection. This is the first analytical closed-form approximation for arbitrary migration rate to appear in the literature. We find that the establishment probability lies between the weak and the strong migration limits if we condition the origin of the mutation to the deme where it is advantageous. This is not the case when we condition the mutation to first occur in a deme where it is disadvantageous. In this case we find that an intermediate migration rate maximizes the probability of establishment. We extend our results to the cases of multiple demes, two demes with asymmetric rates of gene flow, and asymmetric carrying capacities. The latter case allows us to illustrate how density regulation can affect establishment probabilities. Finally we use our results to investigate the role of gene flow on the rate of local adaptation and identify cases in which intermediate amounts of gene flow facilitate the rate of local adaptation as compared to two populations without gene flow.

## Introduction

Studying the maintenance of genetic variation under migration-selection balance has a long tradition in population genetics. While most theoretical research on the establishment and maintenance of local adaptation and population divergence has focused on deterministic models (reviewed in [Felsenstein, 1976, Karlin, 1982, Lenormand, 2002, Nagylaki and Lou, 2008]; see also [Nagylaki and Lou, 2007, Star et al., 2007, Bürger, 009a, Bürger, 009b, Nagylaki, 2009]), considerably less work has been done on the probability of establishment of locally adapted mutations. Even in infinitely large populations, new beneficial mutations experience genetic drift while they are rare, and hence can get lost from the population despite their selective advantage. The probability that a new beneficial mutation evades extinction due to stochastic fluctuations has been called the invasion probability, establishment probability or fixation probability, depending on the context. In the simplest case of a single panmictic population of infinite size, Haldane’s classical result states that the establishment probability of a mutation with time- and frequency-independent selection coefficient s is approximately 2s [Haldane, 1927]. Since then, Haldane’s result has been generalized and extended to several scenarios (see [Patwa and Wahl, 2008] for a review about fixation probabilities of beneficial mutations).

Traditionally, there are two main approaches to study establishment probabilities: branching processes and diffusion approximations. Branching processes often allow for the derivation of simple and intuitive results [Harris, 2002], but are restricted to beneficial mutations in (infinitely large) populations. The diffusion approximation, first used by [Kimura, 1962] in this context, is a powerful tool that allows the derivation of results for both beneficial or deleterious mutations of arbitrary initial frequency in finite populations. The downside is that the derivation of closed form solutions is often harder as compared to branching process, and that the underlying assumptions are not always clear (e.g., the diffusion approximation requires that the mean and variance of the intergenerational change in allele frequencies are small, which is sometimes difficult to interpret in biological terms). Applications of establishment or fixation probabilities include the quantification of the rate of adaption of populations [Orr and Otto, 1994, Wilke, 2004, Desai and Fisher, 2007, Gonçalves et al., 2007], extinction risk due to the accumulation of deleterious mutations [Lynch and Gabriel, 1990], or the rate of emergence of drug resistance or evolutionary rescue [Carlson et al., 2014].

In the context of spatially structured populations, [Barton, 1987] extended the diffusion approximation to account for spatial variation in fitness along a one-dimensional habitat and derived analytical solutions for some special cases. [Kirkpatrick and Peischl, 2013] used a similar approach to study the contribution of new mutations to evolutionary rescue in environments that change in space and time. [McPeek and Holt, 1992] studied the conditions at which a genotype can invade populations fixed for another genotype in environments varying spatially or temporally in a two-patch model. [Tachida and Iizuka, 1991, Gavrilets and Gibson, 2002] and [Whitlock and Gomulkiewicz, 2005] have explored the probability of a single mutant allele fixing in both patches of a two-patch model using diffusion approximations. gavrilets2002Fixation and Whitlock2005probability present the fixation probability as the solution of a system of two quadratic equations that can be solved numerically but so far no closed form solution for the fixation probability has been derived. Building on the results by [Gavrilets and Gibson, 2002], [Wei et al., 2015] derived an explicit formula describing the effect of population size, migration rate, and selection intenisty on the rate of adapative substituions, and identified conditions under which the rate of substitution increases (or decreases) with increasing population size. [Yeaman and Otto, 2011] have used a heuristic ‘‘splicing” approach in which they combine the leading eigenvalues of the transition matrix of a deterministic two-deme model with Kimura’s classical fixation probability formula. Their approach is surprisingly accurate and allowed them to determine the probability of a locally beneficial mutation becoming permanently established and the critical threshold migration rate above which the maintenance of polymorphism is unlikely in finite populations. [Vuilleumier et al., 2008] studied the fixation of locally beneficial alleles through simulations of a metapopulation in a spatially heterogeneous environment. Their findings suggest that a mutation experiencing strong positive selection in parts of an otherwise neutral environment has a higher chance of reaching fixation than a unconditionally beneficial mutation with the same average selection coefficient. This illustrates that heterogeneity in selection coefficients across space can have a large impact on the probability of fixation. Recently, [Aeschbacher and Bürger, 2014] combined multi-type branching processes and diffusion approximations to analytically study the effect of linkage on establishment of locally beneficial mutations in a continent-island model (see also [Yeaman et al., 2016]).

Despite these advances, several open questions remain. Perhaps most importantly, no closed form approximation for the establishment probability in a spatially structured population is available, even for the simplest cases of two-demes with heterogeneous selection (apart from the heuristic formula obtained in [Yeaman and Otto, 2011]). Furthermore, [Vuilleumier et al., 2010] showed that details on how migration is modeled can have a large influence on the outcome on the effects of spatial structure on establishing locally adapted mutations. The same study also identified cases where fixation probabilities lie outside of the range set by low- and high- migration limits, in contrast to what is observed in simpler analytical models [Tachida and Iizuka, 1991, Gavrilets and Gibson, 2002, Whitlock and Gomulkiewicz, 2005].

Deriving a closed expression for the establishment probability in patchy environments is necessary to better understand the role of habitat fragmentation and dispersal on adaptation in spatially heterogeneous environments, for instance in models of evolutionary rescue or evolution of drug resistance [Gomulkiewicz and Holt, 1995, Uecker et al., 2013], studying the spatial origin of mutations that cause range expansion [Behrman and Kirkpatrick, 2011], or studying the role of gene flow on the establishment of local adaptation [Seehausen, 2004].

Here we study the establishment of locally adapted mutations in a discrete migration-selection model using the framework of multi-type branching processes. We present a surprisingly simple and intuitive analytical approximation for the probability that new mutations escape genetic drift and become permanently established. Our results are valid in the limit of weak positive selection in one of the selective habitats but allow for arbitrary migration rates or arbitrarily strong negative selection in the other habitat. We recover previous results for special cases such as weak or strong migration. Our results allow us to quantify the effects of migration on the fate of mutations – depending on whether mutations first occur in individuals living in the deme where the mutation is adapted to or in the deme where the mutations is selected against. We apply our results to biologically interesting scenarios and derive simple results for the effect of asymmetric carrying capacities and density regulation, as well as asymmetric migration, the rate of local adaptation and the contribution of different demes to local adaptation. In particular, we derive conditions under which gene flow between demes facilitates the rate of establishment of locally adapted alleles as compared to the case without gene flow. The latter result is in contrast with common wisdom that gene flow tends to hinder local adaptation, and could have interesting implications for the the role of hybridization during adaptive radiations.

## Results

We start with a symmetric two-deme model to present our model and derive our main result. We then generalize our results to asymmetric migration, multiple demes and account for the effects of density regulation and unequal carrying capacities.

### Two-deme model

We consider an infinitely large population with discrete and non-overlapping generations. The (biologically unrealistic) assumption of an infinitively large population allows us to use branching processes to describe the fate of initially rare mutations, which has been shown to yield accurate results in large but finite populations (e.g., in single panmictic populations the approximation is accurate if *Ns* ≫ 1 where N is the population size and s the selection coefficient of a beneficial mutation [Patwa and Wahl, 2008]). The population is structured into two demes that exchange migrants. At the focal locus, a resident allele is fixed in both demes and a new mutation occurs in a single individual. Each copy of the mutant allele in the population produces a random number of descendant copies in the following generation (“offspring”) that is independent of the number produced by other mutant copies. The mean number of offspring of a mutant copy is given by 1 + *s_i_* in deme *i*x.

In the following, we will focus on the case where selection is acting in opposing directions in the two demes, that is sign(*s*_1_) ≠ sign(*s*_2_), but note that our derivations do not require this assumption. For the remainder we set *s*_1_ > 0 > *s*_2_ without loss of generality. In a haploid population, *s_i_* is the relative fitness advantage (when *s_i_* > 0) or disadvantage (when *s_i_* < 0) of a mutant in deme *i*, while in a randomly mating diploid population, it is the relative fitness (dis)advantage of a heterozygote. Because the ultimate fate of the mutation is decided while mutant homozygotes are still rare, we can ignore their fitness. After reproduction each mutant copy migrates to the other deme with probability *m/2* and remains in its current deme with probability (1 – *m*/2). Thus, *m* = 0 corresponds to two demes without gene flow and *m* = 1 is a special case of the Levene model [Levene, 1953] where allele frequencies are equal in the two patches due to complete mixing of the gene pool every generation. The case *m* = 1 can hence be considered as a single panmictic population with effectively frequency-dependent selection over the whole environment (see also [Ayala and Campbell, 1974]). There are two possible outcomes to the above described process: the mutant allele dies out or it becomes established permanently. Note that establishment does not necessarily imply fixation in both demes in our model: alleles may become permanently established in a balanced polymorphism if migration is sufficiently weak relative to the strength of negative selection against locally disadvantageous alleles. The exact conditions for a balanced polymorphism have been studied extensively in deterministic models, see e.g. [Bulmer, 1972, Nagylaki, 2009, Gavrilets and Gibson, 2002]. We note that a balanced polymorphism may get lost due to drift in finite populations [Yeaman and Otto, 2011] and such a temporary polymorphism is considered as establishment here. We denote by *p*^(*i*)^ the probability of establishment of a mutation that initially appears in an individual in deme *i*.

We model the evolution of the number of copies of a mutant allele that first appears in a single individual using a branching processes with two types of individuals. The type *i* (*i* ∈ {1, 2}) corresponds to the deme in which an individual carrying a copy of the mutant allele resides. We assume that the number of offspring of each copy of the mutant allele is independent of the number of offspring of the rest of the population. The number of mutant copies present in generation *n* can then be described by a vector **X**(*n*) = (*X*_1_(*n*), *X*_2_(*n*)), where *X_i_*(*n*) denotes the number of mutant copies in deme *i* at generation *n* (see Supplemental Material, equation S4). The theory of multi-type branching processes (e.g., [Harris, 2002]) tells us that the vector of extinction probabilities (1 — *p*^(1)^, 1 – *p*^(2)^) is given by

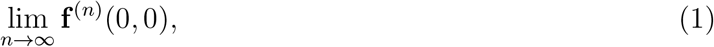

where **f** is the vector of probability-generating functions of the offspring distribution and **f**^(*n*)^ denotes the *n*–fold application of **f**. It can be shown that the establishment probabilities are given by the smallest positive solution of (1– *p*^(1)^, 1– *p*^(2)^) = **f**(1– *p*^(1)^, 1– *p*^(2)^) (see Supplemental Material, equations S1-S6, for details).

Under a Wright-Fisher model of selection and migration, offspring numbers are determined via binomial sampling from the parental generation, which can be approximated by Poisson-distributed offspring in large populations. The mean number of offspring for each type are summarized in table 1 (see Supplemental Material, equations S12-S16 for the derivation of the expected number of offspring per individual). For example, an individual in deme 1 (type 1) will migrate to deme 2 (and hence become a type 2 individual) with probability *m*/2 and then leave an average number of 1 + *s*_2_ offspring. Alternatively, it remains in deme 1 with probability 1 — *m*/2 (and hence remain a type 1 individual) and then leave 1 + *s*_1_ offspring on average. Thus, the expected number individuals of type 1 and 2, respectively, produced by an parent of type 1 will be (1 + *s*_1_)(1 – *m*/2) and (1 + *s*_1_)*m*/2, respectively (see table 1). We assumed here that juveniles migrate before reproduction (and selection) but note that the order of reproduction and migration in the life-cycle does not affect our results.

**Table 1:**
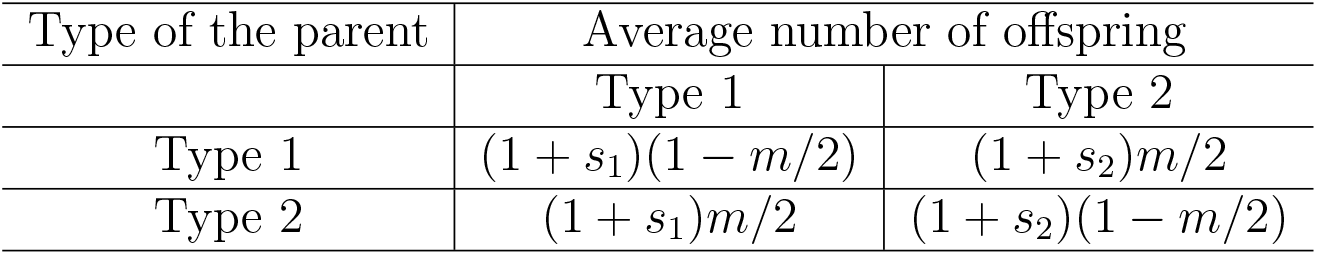
The mean number of individual of type 1 and 2 for parents of type 1 and 2

Since we approximate the binomial sampling as a Poisson distributed offspring, the establishment probabilities are then given by the smallest positive solution of

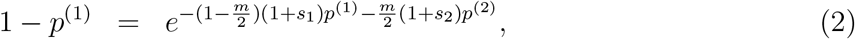

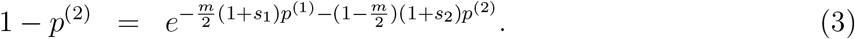

Equations (2) and (3) are transcendental equations for which one can in general not obtain exact solutions and we resort to approximation. We first introduce new parameters that describe the strength of the evolutionary forces relative to the strength of selection for the mutant allele in deme 1: *ζ* = *s*_2_/*s*_1_ and *χ* = *m/s*_1_. Assuming *s*_1_ > 0 > *s*_2_, and taking the limit of weak selection, that is ignoring second- and higher-order terms in *s*_1_, in the Supplemental Material (derivation of equations S24) we show that the establishment probabilities can be written as

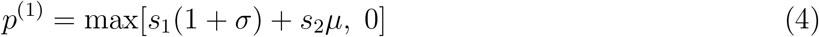

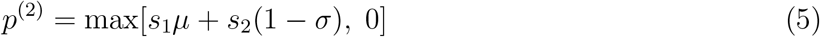

where

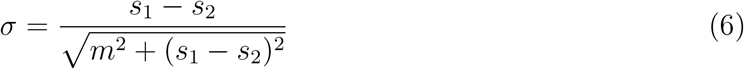

and

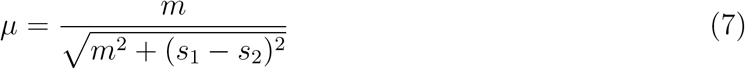

are scaled measures of the heterogeneity in selection and the migration rate, respectively. We note that *σ, μ ∈* [0,1]. Equations (4) and (5) show that the probability of establishment can be written as a weighted sum of the strength of selection in the two demes. Figure 1 shows establishment probabilities for various combinations of selection intensities.

**Figure 1:**
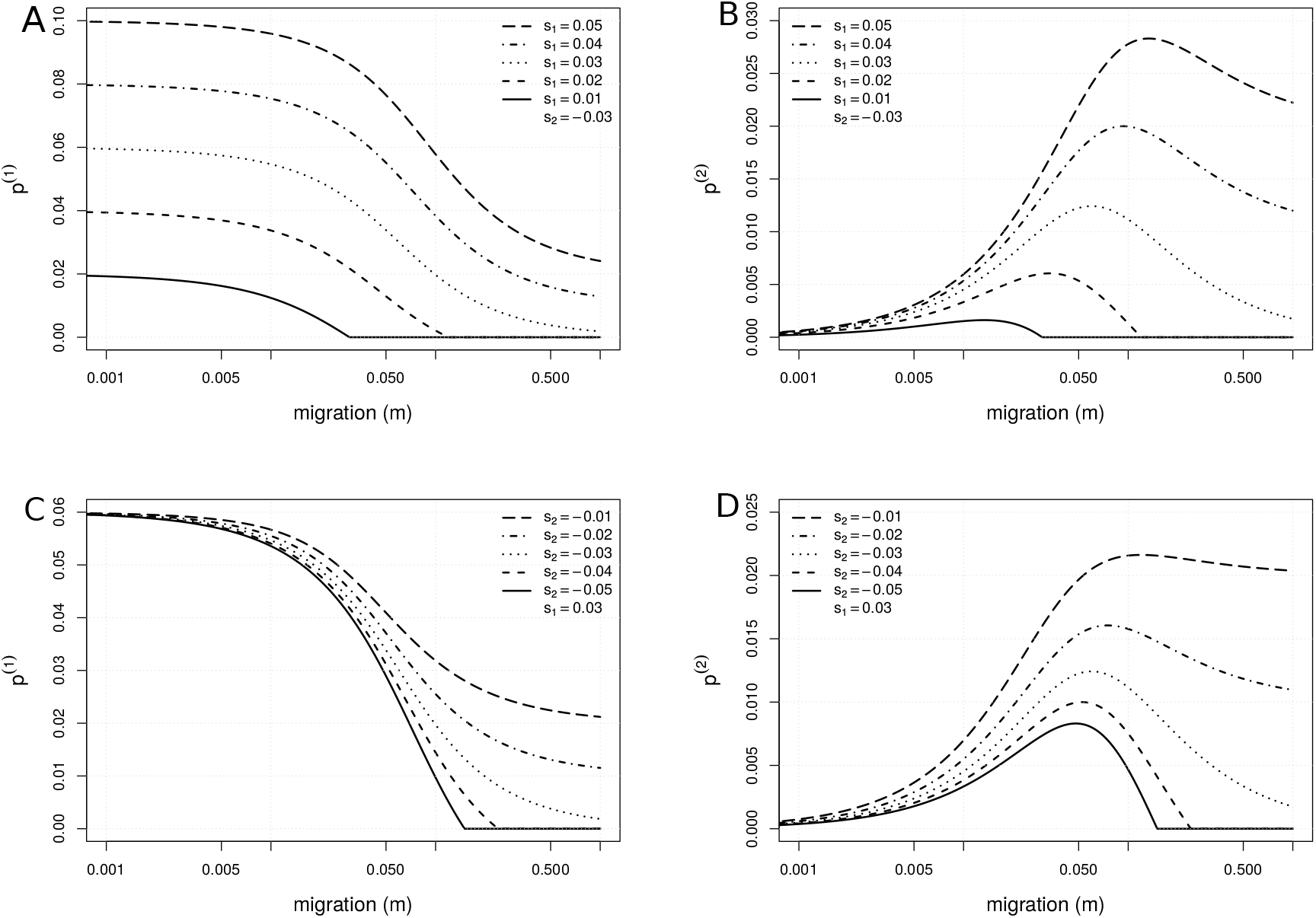
Establishment probabilities as a function of migration rate for various combinations of selection intensities. (A) *p*^(1)^ with fixed *s*_2_ = –0.03. (B) *p*^(2)^ with fixed *s*_2_ = –0.03. (C) *p*^(1)^ with fixed *s*_1_ = 0.03. (D) *p*^(2)^ with fixed *s*_1_ = 0.03.

The weak selection approximation requires that the establishment probability is small but positive (that is, the branching process is slightly supercritical, see [Haccou et al., 2005]). Analysis of the leading eigenvalue of the mean reproduction matrix (S20 and S22 in the Supplemental Material) reveals that the branching process is slightly supercritical if positive selection is weak (*s*_1_ ≪ 1) and either negative selection (*s*_2_) or migration (*m*), but not necessarily both, are sufficiently weak. This implies that our approximation is valid if the mutation is, on average, (strongly) deleterious (*i.e*., *s*_1_ + *s*_2_ < 0) as long as migration is sufficiently weak. Furthermore, if selection is sufficiently weak in both demes (|*s_i_*| ≪ 1), our approximation holds for arbitrarily strong migration. This is sensible because if both migration rate and the strength of negative selection (*s*_2_) are large, mutations will go extinct almost surely and the establishment probability will be 0 (that is, the branching process is subcritical). Figure 2 shows comparison between the analytical approximation and exact solutions of equations (2) and (3), obtained by numerical iteration of the probability-generating function. We find that the approximation shown in (4) and (5) is very accurate with respect to the solutions of equations (2) and (3).

**Figure 2:**
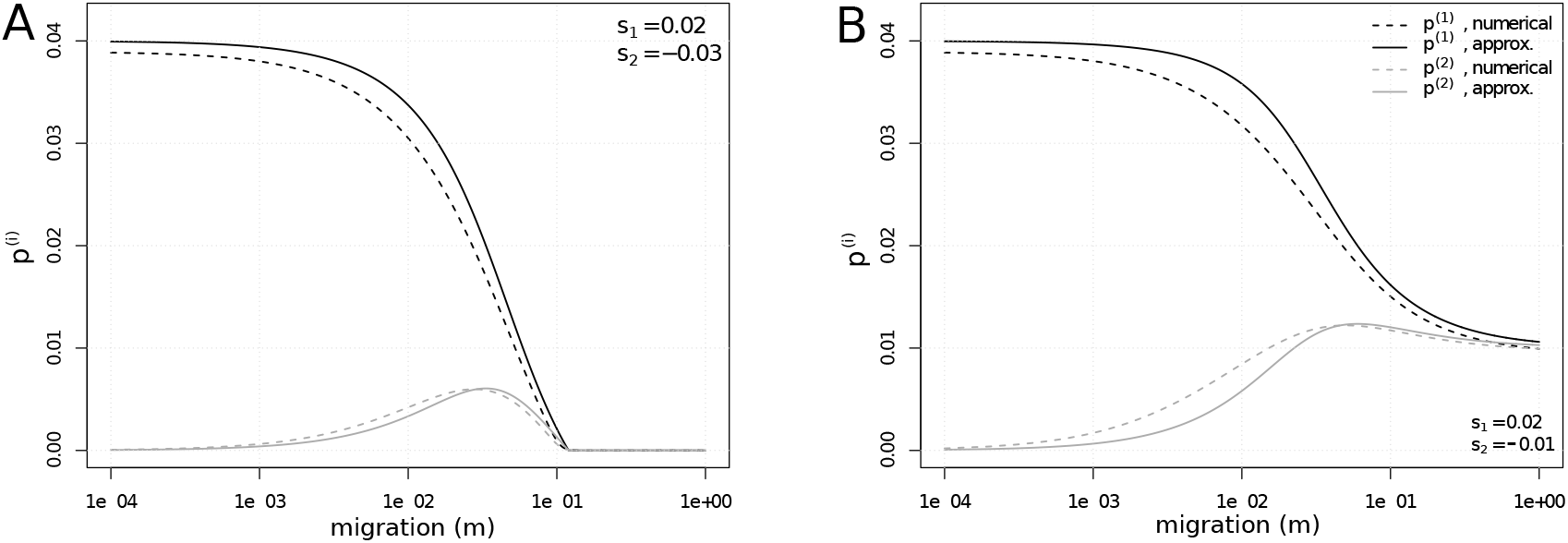
Comparison between exact solution of (2) and (3) and our approximation from equations (4) and (5). The exact solution is obtained numerically after 10’000 iterations of (2) and (3) (see equation (1)). (A) Probabilities of establishment for a scenario where *s*_1_ + *s*_2_ < 0. (B) Probabilities of establishment for a scenario where *s*_1_ + *s*_2_ > 0. The limit for very high migration is *p*^(1)^ = *p*^(2)^ = *s*_1_ + *s*_2_ (see main text).

The establishment probabilities are positive if 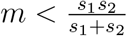 or if *s*_1_ + *s*_2_ > 0. Note that this condition is equivalent to the invasion conditions derived in deterministic models [Bulmer, 1972]. Because *σ* is monotonically decreasing in *m* and *μ* is monotonically increasing in m it follows immediately that *p*^(1)^ is monotonically decreasing in m (see figure 1). For *p*^(2)^ the dependence in m is more complicated. If *m* = 0, it is clear that *p*^(2)^ = 0 because *s*_2_ < 0. Because *p*^(2)^ > 0 if 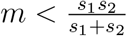 or *s*_1_ + *s*_2_ > 0, *p*^(2)^ is always maximized for some positive migration rate if *s*_1_ > 0 > *s*_2_ *s*_1_+*s*_2_ (figure 1). Straightforward calculations yield that either the maximum of *p*^(2)^ is attained at 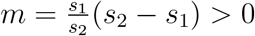 or *p*^(2)^ is monotonically increasing in *m*.

### Comparison with previous results

Equations (2) and (3) recover several previous results for establishment probabilities. In the absence of migration, we have that *σ* =1 and *μ* = 0 and we get *p*^(*i*)^ = max[2*s_i_*, 0], in agreement with Haldane’s classical result for a single panmictic population [Haldane, 1927]. In the limit of strong migration, we get *p*^(1)^ = *p*^(2)^ = max[*s*_1_ + *s*_2_, 0] which means that the establishment probability is determined by the average selection coefficient across demes [Nagylaki, 1980]. In the limit of weak migration, we get 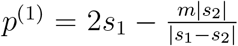 and 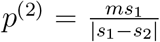. [Gavrilets and Gibson, 2002] used a diffusion approximation to compute fixation probabilities in a biallelic one-locus two-deme model similar to ours. The key difference between our and their approach is that we calculate establishment rather than fixation probabilities, which makes it hard to directly compare our results. Furthermore, no closed-form solution is available for their model. However, in the case where establishment implies fixation, our results are in very good agreement (see Supplemental Material, figure S1). [Yeaman and Otto, 2011] extended classical deterministic two-deme models for diploid individuals (such as [Bulmer, 1972]) to derive a heuristic approximation for the establishment probability of new mutations. They calculate the rate of increase in frequency of a rare locally beneficial mutation and use this initial growth rate as a the selection coefficient in Kimura’s classical equation for fixation probabilities in finite populations [Kimura, 1962]. Numerical comparison reveals a good fit between their results and our equations (4) and (5) (see Supplemental Material, figure S2).

### Asymmetric migration

We next assume that gene flow is asymmetric and let *m_ij_* denote the rate of migration from deme *i* to deme *j*. This includes symmetric migration as a special case if we chose *m_ij_* = *m*/2, for *i* ≠ *j*. In the Supplemental Material (equations S24), we show that the establishment probabilities are then given by

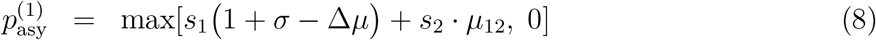

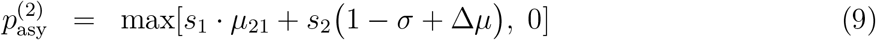

where 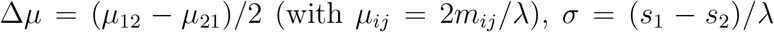. In these last definitions, we used 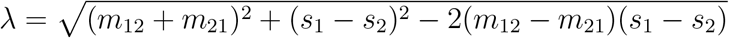. An *m*_crit_ > 0 always exists, such that 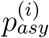 is positive if *m* ∈ [0,*m*_crit_]. Furthermore, 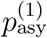 is monotonically decreasing in *m* (see Supplemental Material, eqs. S26-S28). The migration rate for which 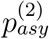 attains its maximum can be easily calculated (see Supplemental Material, equation (S25)). We can see from equations (8)–(9) that selection for locally adapted mutations is amplified as compared to symmetric migration if deme 1 acts as a sink population (*m*_21_ > *m*_12_, and Δ*μ* < 0), and the chances of establishment generally increase with respect to the symmetric migration model (Figure 3). Conversely, if deme 1 acts as a source (*m*_21_ < *m*_12_, and Δ*μ* > 0), selection for locally adapated mutations will be hampered as compared to symmetric migration.

**Figure 3:**
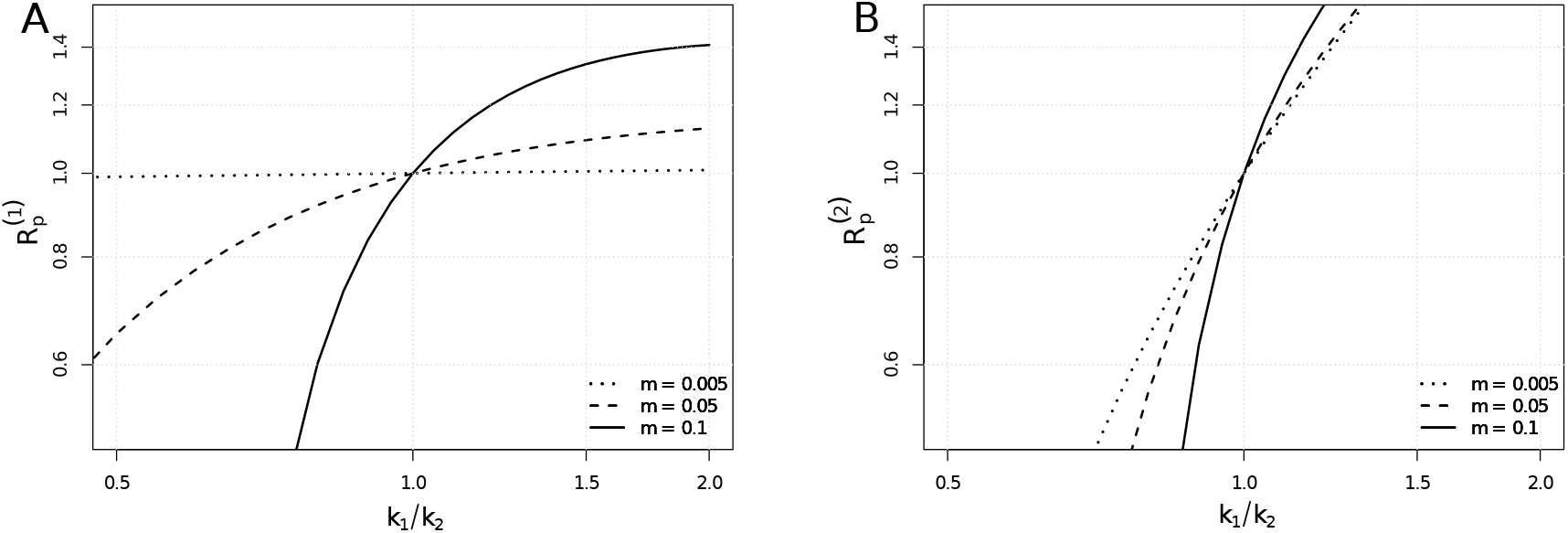
Ratio between probabilities of establishment computed within the model with symmetric migration or within the island model, defined as 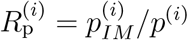, as a function of *k*_1_/*k*_2_. (A) *i* =1 and (B) *i* = 2. In both cases *s*_1_ = 0.02 and *s*_2_ = –0.03.

### Island model with multiple demes

We can extend our results to an island model with multiple demes and two selective habitats. Let s1 and s2 denote the selection coefficients of the mutation in habitat 1 and 2, respectively. We assume that migration occurs at rate m between all demes (i.e., the island model [Wright, 1931]). This model can readily be reduced to a two-deme model with asymmetric migration rates [Whitlock and Gomulkiewicz, 2005]. We assume that all demes are of the same size and that selective habitats 1 and 2 contain *k*_1_ and *k*_2_ = *n* – *k*_1_ demes, respectively. The migration rates between the two selective habitats are then given by *m*_12_ = *mk*_2_/*n* and *m*_21_ = *m*(*n* – *k*_2_)/*n* and the establishment probabilities are given by eqs. (8)–(9) (we call the probabilities of establishment 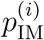 accordingly). If there are more demes where the mutation is beneficial (habitat 1) than where the mutation is detrimental (habitat 2), we have more individuals migrating from habitat 2 to habitat 1 and hence Δ*μ* < 0. As a consequence the contribution of selection in deme 1 is amplified and establishment probabilities are generally larger as compared to the case with two equally large habitats (see eqs. (8) and (9), see also figure 3).

### Asymmetric carrying capacities and density regulation

So far we ignored the effects of density regulation because we assumed infinitely large populations. We next modify the offspring distribution in our branching process to account for the effects of deme-independent density regulation (soft selection, sensu [Wallace, 1975]) in a model with symmetric migration. Let *κ*_1_ and *κ*_2_ denote the carrying capacities of deme 1 and 2, respectively. The larger deme then acts as a source, that is, it sends out more migrants than it receives. Here we assume that density regulation acts after migration and brings each deme back to its carrying capacity instantaneously. Initially both demes are at carrying capacity. The number of individuals in deme *i* after migration but before density regulation are denoted 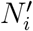 and are given by

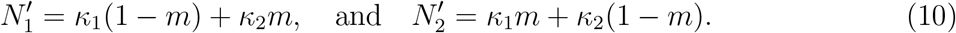

Density regulation will then change the number of individuals in each deme by a factor

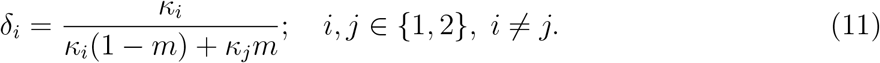

We can integrate this effect of density regulation in our branching process framework by modifying the absolute fitness of individuals in deme *i* to *w_i_* = (1 + *s_i_*)*δ_i_* (see equation (S32)).

The establishment probabilities for this case are explicitly calculated in the Supplemental Material (see equations (S33)).

If *κ*_1_ < *κ*_2_, deme 1 receives more migrants than it sends out and density regulation will cull the population size back to carrying capacity. Thus, in a certain sense deme 1 is behaving like a shrinking population, which should reduce the establishment probability of mutations that are beneficial in that deme [Otto and Whitlock, 1997]. Our results confirm this intuition (figure 4) and show that the establishment probability increases if the deme where the allele is beneficial has a smaller carrying capacity. Likewise, if *κ*_2_ < *κ*_1_, deme 1 is growing after migration, which reduces drift and increases the establishment probability of mutations that are adapted to deme.

**Figure 4:**
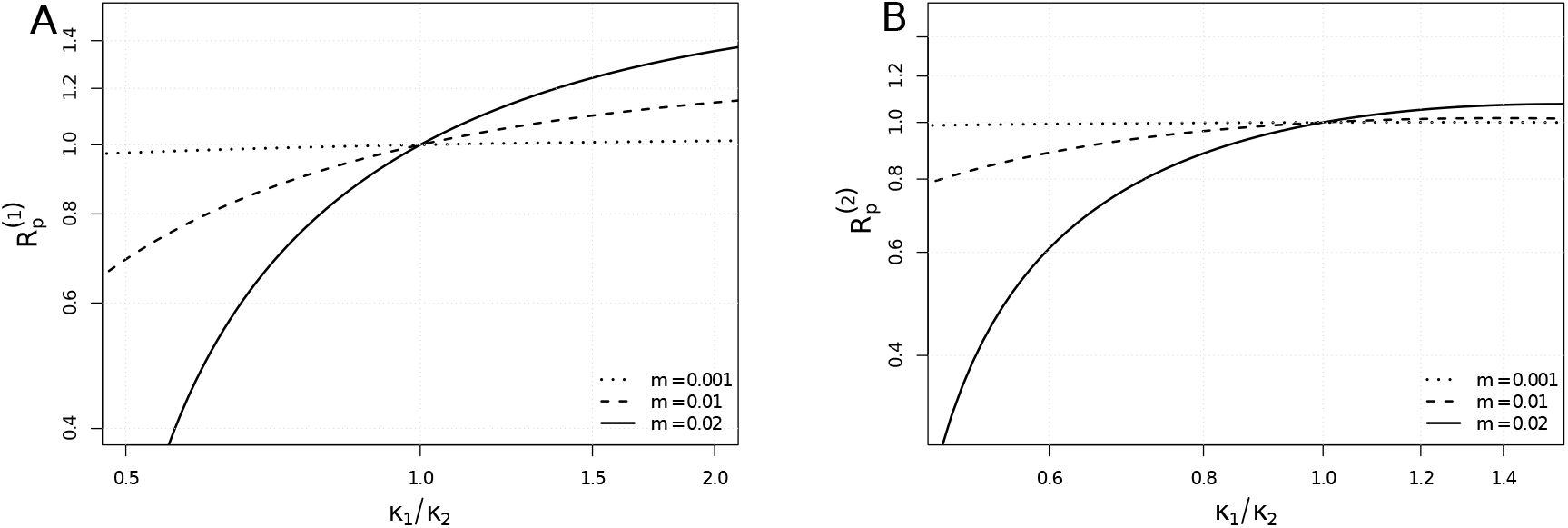
Ratio between probabilities of establishment computed for the model with symmetric migration and for the model with asymmetric carrying capacities (see equations for 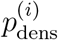, Supplemental Material, equation (S33)), defined as 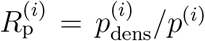, as a function of *κ*_1_/*κ*_2_. (A) *i* =1 and (B) *i* = 2. In both cases *s*_1_ = 0.02 and *s*_2_ = –0.03.

### Comparison with simulations

In single, panmictic populations, the branching process assumption requires that the population is sufficiently large relative to the strength of (positive) selection [Patwa and Wahl, 2008]. While the applicability of multi-type branching processes to study establishment probabilities has been established in previous studies [Haccou et al., 2005], the exact conditions under which the branching process approximation is expected to be accurate in structured populations of finite size remain unclear and are difficult to derive (especially in the absence of solutions for models with finite population size for comparison). To asses the quality of our approximation in finite populations we therefore resort to comparison with simulations for small populations (*N* = 300 or 400 individuals) and moderately strong positive selection (*s*_1_ = 0.02) such that *Ns*_1_ > 1. Results from simulations show a good fit with analytical results (see figure 5) for the case of symmetric demes, asymmetric migration or asymmetric carrying capacities. Simulations were performed assuming logistic density regulation [Beverton and Holt, 1957], and inserting one mutant in either deme 1 or 2 and letting the system evolve for 20000 generations. 50000 replications are done for each simulated scenario. Our approximation tends to overestimate *p*^(1)^ and underestimate *p*^(2)^. This is expected, since the same behavior can be seen when we compare the approximation to the exact solution (see figure 2).

**Figure 5:**
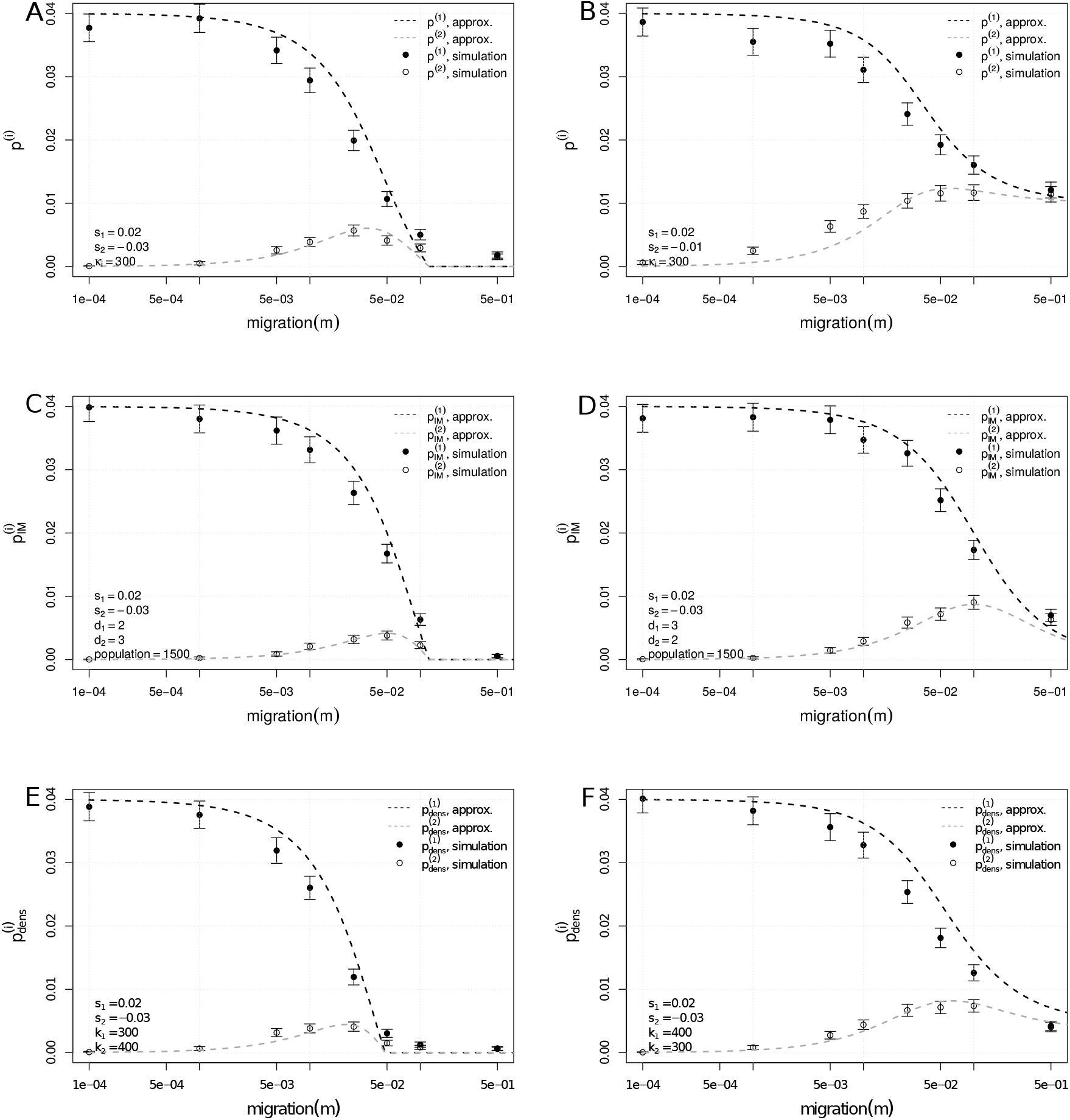
Comparison of simulations and analytical approximations. Model with symmetric migration and symmetric demes with *s*_1_ = 0.02, and *s*_2_ = –0.03 (A), and *s*_1_ = 0.02, and *s*_2_ = –0.01 (B). Island model with two demes of type 1 (*d*_1_ = 2, *s*_1_ = 0.02) and three demes of type 2 (*d*_2_ = 3, *s*_2_ = –0.03) (C), and with three demes of type 1 (*d*_1_ = 3, *s*_1_ = 0.02) and two demes of type 2 (*d*_2_ = 2, *s*_2_ = –0.03) (D). Model with different carrying capacities, with carrying capacities *κ*_1_ = 300 and *κ*_2_ = 400 (E), and *κ*_1_ = 400 and *κ*_2_ = 300 (F). The analytic formula for 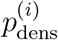, shown in images (E) and (F), is described in the Supplementary Information (equations S33).

### Global rate of establishment of locally adapted alleles

It is commonly assumed that gene flow hampers or even prevents local adaptation [Lenormand, 2002]. Here we use our results to quantify the effect of migration on the overall establishment probability, defined as

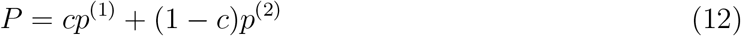

where *c* and 1 – *c* denote the relative sizes of deme 1 and deme 2, respectively.

We next take the derivative of *P* with respect to *m* at *m* = 0. If this derivative is positive, the rate of adaptation increases when we introduce some gene flow as compared to the case without gene flow between demes. In other words, we derived the condition under which the unconditional establishment probability of locally adapted mutations increases when we introduce small amounts of migration. We find that this is the case in all our models if

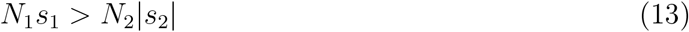

where *N*_1_ and *N*_2_ are the number of individuals in deme 1 and 2 respectively. Hence, in the symmetric model where the probability of establishment is given by (4) and (5), we find that gene flow increases the chances of establishment when |*s*1| > |*s*_2_|. Therefore, if the selective advantage in one deme is larger than the selective disadvantage in the other deme, some gene flow can facilitate the establishment of local adaptation as compared to completely isolated demes. This can also be seen directly via the weak migration approximation 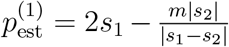 and 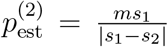 derived above. If we consider the model with asymmetric migration, the condition becomes

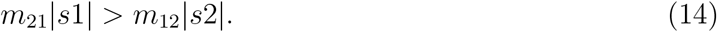

Using the definitions of *m*_12_ and *m*_21_ derived for an multi-deme island model, equations (13) and (14) are identical.

## Discussion

We used multi-type branching processes to study the establishment of locally beneficial mutations in a spatially heterogeneous environment with two selective habitats. Our main result is a simple and analytical closed-form approximation for the probability of establishment of a locally beneficial mutation in a two-deme model with divergent selection and symmetric migration between demes (equations (4) and (5)). By establishment we mean that a mutation permanently establishes in the meta-population, either by going to fixation or by maintenance as a balanced polymorphism. To our knowledge this is the first closed-form analytical approximation for an establishment probability in this context that is valid for arbitrary migration rates (but see [Yeaman and Otto, 2011] for a heuristic approach). The resulting formula is intriguingly simple and intuitive: the probability of establishment is simply a weighted average over selection coefficients in the two demes, where the weights are determined by the relative contributions of migration and spatial variation in selection. We extended our main result to asymmetric migration between two demes, a multi-deme island model with two selective habitats, and studied the impact of variation in carrying capacities and density regulation on the establishment of locally adapted mutations. We show that establishment probabilities can fall outside the range spanned by the weak or strong migration limits, and provide conditions for when this is the case. In particular, we identify conditions under which small amounts of migration can facilitate the build-up of adaptive divergence as compared to two demes without gene flow. Examination of the weak selection approximation provides an intuitive explanation for this phenomenon. On the one hand, migration removes mutations from the deme where they adapted to and hence decreases the establishment probability in deme 1 from 2*s*_1_ to 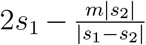. On the other hand, gene flow allows mutations occurring in the wrong habitat, which are doomed to extinction in the absence of gene flow, to eventually reach the right habitat and then spread there. Thus, gene flow increases the establishment probability from 0 to 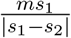 for mutations occurring in a deme in which they are maladaptive. Taken together the net effect of migration will therefore be positive if positive selection in one deme is sufficiently strong relative to negative selection in the other deme. For instance, if the two demes are of equal size, this is the case if |*s*_1_| > |*s*_2_|. Our derivation assumes an (infinitely) large population, that selection for the beneficial mutation is sufficiently weak, and that either migration or negative selection is not too large. With respect to finite population sizes, we expect that our approximation should hold in finite populations as long as the product of population size and selection coefficient s1 is larger than 1 (*Ns*_1_ ≫ 1, see fig.5), in analogy to single panmictic populations. However, the exact conditions for when our approximation is accurate will depend on the population sizes in both demes, the strength of positive as well as negative selection, and the amount of gene flow and are difficult to derive and beyond the scope of this study. Worthy of note, our approximation remains valid if the mutation is strongly deleterious (or even lethal) in one of the demes (see figure S3 in the Supplementary Material), and holds for arbitrarily strong migration if negative selection is sufficiently weak. Furthermore, we have modeled a haploid population to avoid the intricacies of dominance. However, our results should remain valid in diploid populations if we use the fitness of the mutant allele in our model as the fitness of heterozygotes in a diploid model. The ultimate fate of a mutation will be determined while it is rare so that we can ignore the fitness of homozygotes.

We have been discussing single mutations in isolation and neglected genetic events that may interfere with the establishment process (e.g., clonal interference [Gerrish and Lenski, 1998]). Our results should therefore hold in sexually reproducing species with strong recombination. Our approximation is less plausible in organisms that reproduce with little or no recombination, such as most microbes, or for mutations in genomic regions with low recombination rates. Competition between simultaneously spreading beneficial mutations (clonal interference) can have severe impacts on each other’s establishment [Gerrish and Lenski, 1998, Orr, 2000]. This effect is difficult to account for [Wilke, 2004] and is most important in populations with little or no recombination, because recombination can break associations between mutations that occur in different parts of a genome [Muller, 1932, Hill and Robertson, 1966]. A second effect that we did not account for is the genomic background on which the mutation falls via additive or epistatic interactions between mutations. This can be sidestepped by interpreting the selection coefficient as referring to the focal individuals fitness rather than to the effect of the mutation. Recently, [Aeschbacher and Bürger, 2014] and [Yeaman et al., 2016] have studied the establishment of islands of differentiation in a continent island model and explicitly modeled two different genetic backgrounds using a multi-type branching process, similar to the approach used here. While it would be very interesting to combine our approach with their analysis, the algebraic challenges of studying a branching process with four different types will make analytical progress challenging.

Previous work based on individual based simulations has shown that variation in densities across demes can affect the establishment of new mutations [Vuilleumier et al., 2010]. Our results confirm this and show that selection is more efficient in source-demes. We show that the combination of density regulation and asymmetric migration mimics the effects of a growing population, which increases the absolute fitness of individuals and leads to more efficient positive selection [Otto and Whitlock, 1997].

The solution that we presented in this paper also assumes that migration rates remain fixed in time. We know, however, that spatially varying selection can lead to evolution of dispersal [Ronce, 2007]. Our model is therefore plausible if migration is mainly determined by geographical features of the environment, or if there is little or no genetic variability for traits related to dispersal. Furthermore, variation in density can introduce non-random movement between demes. We have accounted for this in eqs. (8) and (9) where we allowed migration rates to be asymmetric. In equations (8) and (9), the carrying capacity of each type of habitat is not explicitly taken into account. A combination of the result for asymmetric migration rates with the scenario with density regulation (see equations S33) may be necessary to study extreme cases in which asymmetry of migration is related with asymmetry in carrying capacities. While this should be possible in a multi-type branching process framework, the derivations are complicated and are beyond the scope of this work. An alternative approach would be to model the evolution of dispersal traits deterministically, which would yield a time-dependent multitype branching process that could be studied with a perturbation approach similar to the one derived in [Peischl and Kirkpatrick, 2012].

Our predictions could be tested in experimental metapopulations, for instance, using a setup similar to the one in [Bell and Gonzalez, 2011], where gene flow was mimicked via pipetting in experimental yeast metapopulations. Interestingly, in their experiment, [Bell and Gonzalez, 2011] found that gene flow can facilitate the chance for evolutionary rescue in a gradually deterioration spatially extended population relative to the case of no gene flow, which could be explained by our theoretical predictions (see also [Uecker et al., 2013]). Another promising system to test our predictions experimentally would be the ciliate model organism *Tetrahymena,* where metapopulations can be created as microcosms of connected vials and selection intensities could be controlled with antibiotic concentrations [Altermatt et al., 2015]. Empirical evidence for our predictions will be somewhat harder to obtain, as crucial factors such as the fitness effects of mutations as well as the amount of gene flow are hard to measure.

We have presented here a mathematically rigorous approximation of establishment probabilities in a spatial framework using the theory of multi-type branching processes. It would be very interesting to generalize our approach to more than two different types of individuals. While the theoretical foundation is laid out, finding actual solutions for establishment probabilities in higher-dimensional system poses algebraic challenges that might be difficult to overcome. Nevertheless, the simple and intuitive form of our solution suggests that this approach can be exploited further and that our results can be generalized and extend to various scenarios that include more than two types of individuals.

## Acknowledgements

We thank Reinhard Bürger for stimulating discussions on the subject. We gratefully acknowledge helpful comments from three anonymous reviewers.

